# Evaluation of phenotyping errors on polygenic risk score predictions

**DOI:** 10.1101/724534

**Authors:** Ruowang Li, Jiayi Tong, Rui Duan, Yong Chen, Jason H. Moore

## Abstract

Accurate disease risk prediction is essential in healthcare to provide personalized disease prevention and treatment strategies not only to the patients, but also to the general population. In addition to demographic and environmental factors, advancements in genomic research have revealed that genetics play an important role in determining the susceptibility of diseases. However, for most complex diseases, individual genetic variants are only weakly to moderately associated with the diseases. Thus, they are not clinically informative in determining disease risks. Nevertheless, recent findings suggest that the combined effects from multiple disease-associated variants, or polygenic risk score (PRS), can stratify disease risk similar to that of rare monogenic mutations. The development of polygenic risk score provides a promising tool to evaluate the genetic contribution of disease risk; however, the quality of the risk prediction depends on many contributing factors including the precision of the target phenotypes. In this study, we evaluated the impact of phenotyping errors on the accuracies of PRS risk prediction. We utilized electronic Medical Records and Genomics Network (eMERGE) data to simulate various types of disease phenotypes. For each phenotype, we quantified the impact of phenotyping errors generated from the differential and non-differential mechanism by comparing the prediction accuracies of PRS on the independent testing data. In addition, our results showed that the rate of accuracy degradation depended on both the phenotype and the mechanism of phenotyping error.

## 1. Introduction

Understanding the risk factors underlying diseases has long been pursued in healthcare in order to screen and prevent disease onset for high-risk individuals. Proper quantification of the risk factors could help stratify patients based on their risk profiles, which in turn can be beneficial for developing personalized disease prevention and treatment strategies^1^. With the development of high-throughput sequencing technologies, it is now a reality to systematically evaluate the genetics’ contribution to disease risks. Genetic twin studies have shown that many human phenotypes and diseases are highly hereditable; however, early genome-wide association studies have identified many single nucleotide polymorphisms (SNPs) that are only weakly to moderately associated with the diseases. In addition, for the associated SNPs, they only explain a small amount of the disease risks^2–4^. Recent studies have discovered that many phenotypes are polygenic in nature, meaning a phenotype is associated with more than one gene^5,6^. Thus, the polygenic risk score (PRS) method was developed to capture the small effects from many genetic factors in order to combine their effects into a single predictive variable^6,7^. The PRS has been evaluated for its role in determining disease risk in many complex diseases including coronary artery disease, atrial fibrillation, type 2 diabetes, inflammatory bowel disease, breast cancer^8^, obesity^9^, schizophrenia^10^, and antipsychotic drug treatment^11^. For some of the diseases, the predictive power of PRS has reached clinical significance similar to that of monogenic mutations^8^.

For the past decade, electronic health record (EHR) linked genetic data has proven to be a valuable data source for identifying genetic associations for diseases. EHR with linked genetic data has the advantages of having a large sample of the patient population as well as a rich source of matching clinical phenotypes to conduct genomics research. In addition, several EHR data have already been used to conduct PRS research, including the UK Biobank^8^ and eMERGE^12^. While the genetic data is an integral part of PRS prediction, the phenotype used to construct PRS is equally as important. A crucial step in constructing a PRS is to determine the marginal association of each SNP with the phenotype. Thus, the quality of the associations determines the utility of the constructed PRS. However, there are unavoidable biases and measurement errors associated with the EHR derived phenotypes. Existing studies have evaluated the impact of phenotyping errors on statistical inference and showed that the errors decreased the power^13^ as well as inflated the type 1 error^14^ of the associations. Nevertheless, so far, there has been no investigation on the impact phenotyping error on the predictive ability of PRS.

In this study, we used real EHR data from eMERGE to simulate three types of phenotype under two phenotyping error mechanisms. We systematically quantified the PRS predictive ability in different phenotypes under different severities of phenotyping error and error mechanisms. Our results showed that as more errors were added to the phenotypes, non-differential phenotyping errors lowered the PRS prediction accuracies similarly among different phenotypes. In contrast, differential phenotyping errors affected the PRS prediction differently depending on the underlying phenotype model. We believe that our results could better inform researchers and clinicians of the robustness of PRS when assessing disease risk.

## 2. Method

To evaluate the impact of phenotyping error on PRS prediction, we used simulated datasets where we knew the ground truth to quantify the change in prediction accuracy. The evaluation was carried out in five stages. 1) Use real patients’ genetic data from eMERGE EHR as input to construct PRS. 2) Simulate known phenotypes under various underlying true models. The phenotypes were constructed to have true associations with demographic, environmental, clinical, and genetic factors (PRS). 3) Inject errors into the known phenotypes under two different error generating mechanisms: differential and non-differential 4) Adjust the strength of the phenotyping error 5) Quantitatively evaluate the predictive ability of PRS on the testing data under each simulation scenario.

### 2.1. eMERGE EHR genetic data

In order to simulate realistic PRS, we utilized the patients’ genetic data from the electronic medical records and genomics network (eMERGE, dbGaP accession: phs000888.v1.p1)^15^. Recent studies suggested that PRS does not perform well across multiple ethnic groups; thus we restricted our study samples to only one ethnicity^16,17^. To maximize the sample size, we extracted white patients from nine different hospitals under eMERGE: Children’s Hospital of Pennsylvania, Cincinnati Children’s Hospital Medical Center/Boston’s Children’s Hospital, Geisinger Health System, Group Health/University of Washington, Essentia Institute of Rural Health, Marshfield Clinic, Pennsylvania State University (Marshfield), Mayo Clinic, Icahn School of Medicine at Mount Sinai School, Northwestern University, and Vanderbilt University. The SNP genotyping was performed using the Illumina 660W-Quad BeadChip at the Center for Genotyping and Analysis at the Broad Institute, Cambridge, MA. Genome imputation was performed by eMERGE according to the standard pipeline^18^. Overall, 31,183 patients’ 38,040,165 autosomal SNP genotypes were extracted.

### 2.2. Phenotype Simulation

We simulated three types of phenotype under different underlying true models (Figure 1, solid arrows on top). First, a phenotype was simulated to be associated with the demographic variables, a set of causal SNPs, and an environmental factor. All variables were independently associated with the phenotype; thus, it was named the *independent model.* Second, a phenotype was simulated to be associated with the demographic variables, a set of casual SNPs and a related diagnosis. In this case, the related diagnosis was also associated with a subset of the causal SNPs, though the associations were different from that of the phenotype. For example, a subset of causal SNPs may have pleiotropic effects between hypertension and heart failure, but the pleiotropic associations with the two diseases are distinct. In addition, diagnosis in hypertension is also one of the factors in determining heart failure status. Because the related diagnosis (hypertension) shared a subset of causal SNPs with the phenotype (heart failure) and the associations were distinct, the model was called the *weakly correlated model.* Finally, a phenotype was similarly simulated to be associated with demographic variables, a set of causal SNPs, and a related diagnosis as in the *weakly correlated model.* However, the set of pleiotropic SNPs had the same effects on the related diagnosis as on the phenotype. An example would be that a subset causal SNPs are similarly associated with cardiac arrest (related diagnosis) as well as heart failure (phenotype). Furthermore, cardiac arrest is also associated with heart failure diagnosis. In this study, this model was named *strongly correlated model.* The SNPs in all models were randomly selected from the common SNPs (minor allele frequency > 5%) in the eMERGE EHR genetic data. The mathematical models for the phenotype simulation are presented in the following sections.

**Fig. 1.**
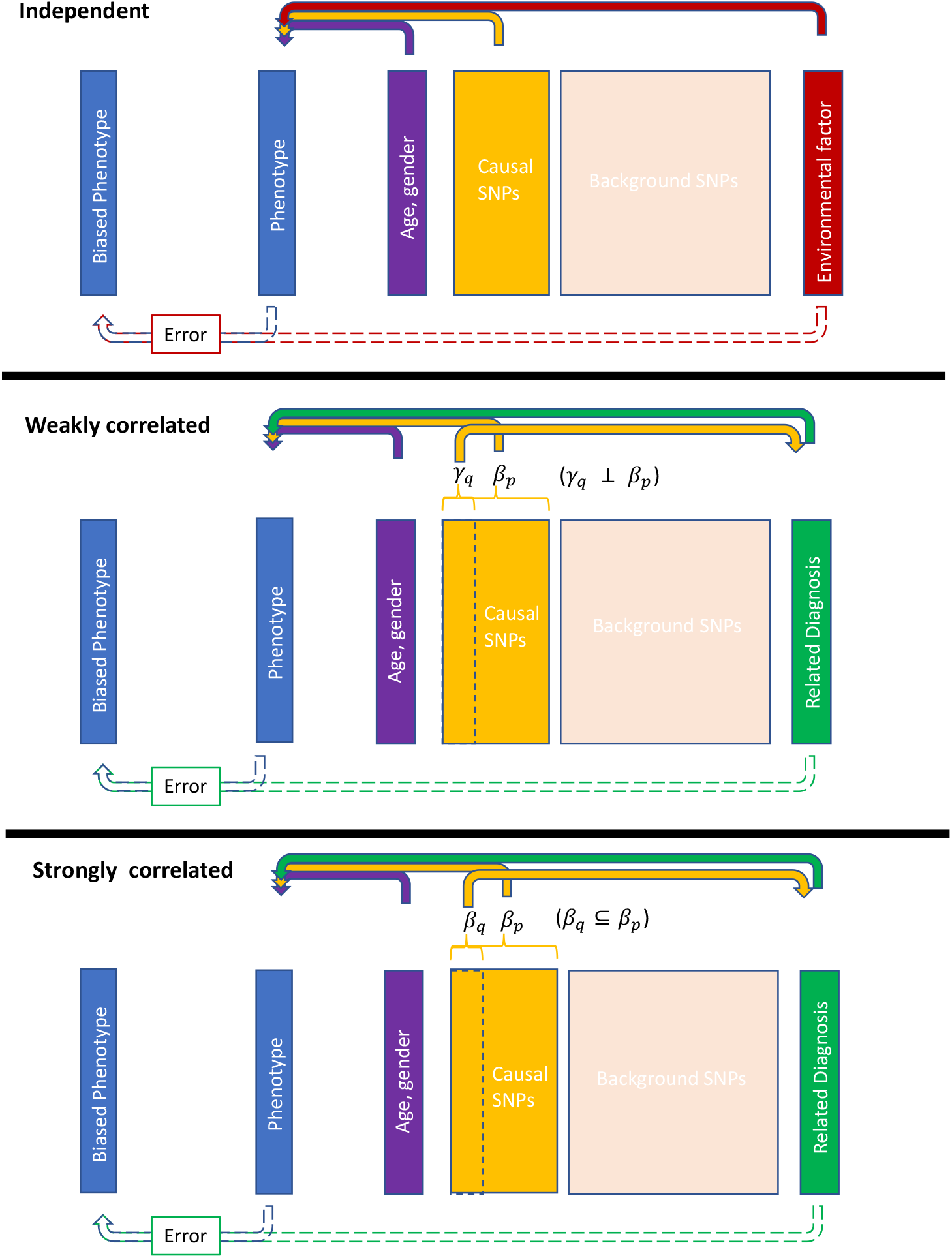
Phenotypes generating mechanism. The phenotypes were generated using patients’ age, gender, SNP genotypes, and an environmental factor or a related diagnosis status. The top solid arrows represent the true phenotype generating mechanism. In the *independent model*, all factors were independently associated with the phenotype. In the weakly correlated model, the related diagnosis and the phenotype shared a subset of causal SNPs, but the associations *γ_p_* and *β_p_* were independent. In the *strongly correlated model*, the subset of shared casual SNPs had the same associations, as in *β_q_* is a subset of *β_p_*. The bottom dotted arrows indicate the phenotype error generating mechanism. The biased phenotypes were generated based on the values of the true phenotype and the environmental factor or the related diagnosis.

#### 2.2.1. Independent model

In this model, the phenotype Y was generated through the logistic model.

Phenotype:

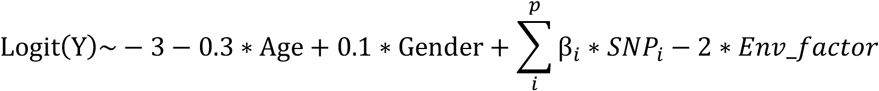

The coefficients for the intercept, age, gender, and environmental factors (Env_factor) were selected so that the disease prevalence was around 30%. The same coefficients were also used for the *weakly correlated* and the *strongly correlated model* so that the models were comparable. The distributions of the random variables in all equations were listed in Table 1.

**Table 1.** Parameter values for phenotype simulation

| Variable | Value |
| --- | --- |
| Total randomly selected SNPs | 500 |
| Phenotype associated SNPs | $p = 100$ |
| Diagnosis associated SNPs | $q = 50$ |
| Age | Normal (40, 10) |
| Gender | Bernoulli ($p = 0.5$) |
| Environmental factor (Env_factor) | Bernoulli ($p = 0.5$) |
| Phenotype ~ SNP associations | $\beta \sim \text{Normal}(0, 0.3)$ |
| Related diagnosis ~ SNP associations | $\gamma \sim \text{Normal}(0, 0.3)$ |

#### 2.2.2. Weakly correlated model

In the *weakly correlated model*, a related diagnosis was first generated using q SNPs, where q was a subset of p SNPs that were used to generate the phenotype. In addition, the coefficients *γ* for generating the related diagnosis were independent of β that were used to generate the phenotype.

Related diagnosis:

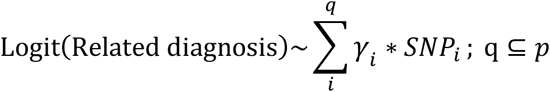

Phenotype:

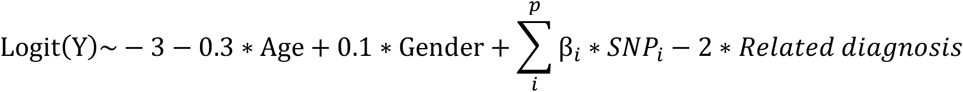

#### 2.2.3. Strongly correlated model

The *strongly correlated model* was the same as the *weakly correlated model* except that the related diagnosis and the phenotype shared a subset of q SNPs as well as their coefficients.

Related diagnosis:

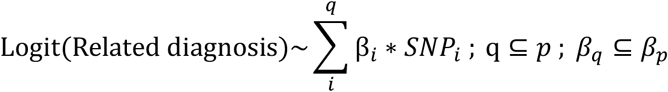

Phenotype:

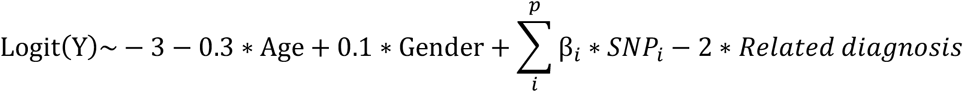

### 2.3. Biased phenotype due to errors

As shown in Figure 1, the biased phenotypes were generated based on the value of the true phenotypes as well as the environmental factor or the related diagnosis (Figure 1, dotted arrows at the bottom). The intuition was that, first, the biased phenotype would be expected to be a deviation from the true phenotype. Second, many of the phenotyping algorithms utilized by EHR systems used environmental and diagnosis variables to determine the phenotype or disease status, thus, the precision of the phenotype was also associated with these factors^19–21^. Mathematically, the phenotyping errors were determined by the sensitivity and specificity:

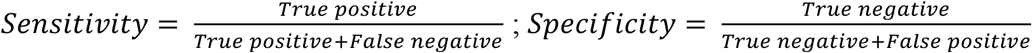

In the independent model, the biased phenotype was generated using the following 2Ô2 tables.

|  | Y=1 | Y=0 |
| --- | --- | --- |
| ENV_FACTOR = 1 | Sensitivity<sub>Exposure</sub> | Specificity<sub>Exposure</sub> |
| ENV_FACTOR = 0 | Sensitivity<sub>non_Exposure</sub> | Specificity<sub>non_Exposure</sub> |

For example, the Sensitivity_Exposure_ controlled the sensitivity of the biased Y when the true phenotype Y =1 and Env_factor = 1. The new phenotype value under this combination was generated using the Bernoulli distribution with the probability equaled to Sensitivity_Exposure_. In contrast, the Specificity_Exposure_ determined the probably of the biased Y = 0, when true Y = 0 and the Env_factor = 0. The value was generated by Bernoulli (1-SpecificityExposure). Thus, the degree of phenotyping errors was controlled by the values of the sensitivity of specificity. As a special case, a phenotype was the gold standard when sensitivity = specificity = 100%.

For biased phenotypes, the phenotyping error was non-differential when the sensitivities and specificities were the same across the two Env_factor levels; otherwise, the error was differential. For instance, a phenotype that is more error-prone for patients with lower levels of environmental exposure would be differentially biased.

NON-DIFFERENTIAL PHENOTYPING ERROR
|  | Y=1 | Y=0 |
| --- | --- | --- |
| ENV_FACTOR = 1 | a% | b% |
| ENV_FACTOR = 0 | a% | b% |

DIFFERENTIAL PHENOTYPING ERROR
|  | Y=1 | Y=0 |
| --- | --- | --- |
| ENV_FACTOR = 1 | a% | c% |
| ENV_FACTOR = 0 | b% | d% |

The biases for the *weakly correlated* and *strongly correlated models* were generated in the same fashion except that the Env_factor was replaced with the related diagnosis.

### 2.4. Biased phenotype generation

For all phenotypes (*independent, weakly correlated*, and *strongly correlated*), a range of phenotyping errors were introduced using different levels of sensitivity and specificity. In addition, differential and non-differential error generating mechanisms were applied at each sensitivity and specificity level. To simplify the presentation of the results, the same value of sensitivity and specificity for the non-differential phenotyping error was used (Table 2). For differential phenotyping error, one sensitivity and specificity were kept at 99%, while the others varied (Table 3). Overall, 60 biased phenotypes were generated.

**Table 2.** Sensitivity and specificity for the non-differential phenotyping error

| Error model name | Error mechanism |  |  |
| --- | --- | --- | --- |
| **Gold standard** | No error |  |  |
| **X = (95, 90, 85, 80, 75, 70, 65, 60, 55, 50)** |  | Y=1 | Y=0 |
|  | Env_factor or Diagnosis =1 | Sensitivity = X% | Specificity = X% |
|  | Env_factor or Diagnosis =0 | Sensitivity = X% | Specificity = X% |

**Table 3.** Sensitivity and specificity for the differential phenotyping error

| Error model name | Error mechanism |  |  |
| --- | --- | --- | --- |
| **Gold standard** | No error |  |  |
| **X = (95, 90, 85, 80, 75, 70, 65, 60, 55, 50)** |  | Y=1 | Y=0 |
|  | Env_factor or Diagnosis =1 | Sensitivity = X% | Specificity = 99% |
|  | Env_factor or Diagnosis =0 | Sensitivity = 99% | Specificity = X% |

### 2.5 Evaluation of PRS prediction

The effect of the phenotyping errors on PRS prediction was evaluated in the following steps.

|  |  |
| --- | --- |
| 1. Split the data into training and testing | $\text{Data} = 70\% \text{Data}_{\text{train}} + 30\% \text{Data}_{\text{test}}$ |
| 2. Sample sizes of the data split | Training: 21,827; Testing: 9355 |
| 3. Obtain coefficients for the SNPs using the training data | $Y_{\text{train}} \sim \hat{\beta}_{i,\text{train}} * \text{SNP}_{i,\text{train}}$ |
| 4. Construct PRSs in both training and testing data | $\text{PRS}_{\text{train},p} = \sum_{i=1}^p \hat{\beta}_{i,\text{train}} * \text{SNP}_{\text{train},i}$ $\text{PRS}_{\text{test},p} = \sum_{i=1}^p \hat{\beta}_{i,\text{train}} * \text{SNP}_{\text{test},i}$ |
| 5. Build a predictive model using the PRS in the training data | $Y_{\text{train}} \sim \hat{\lambda}_{\text{PRS},\text{train}} * \text{PRS}_{\text{train},p}$ |
| 6. Apply the model to the test data | $\hat{Y}_{\text{test}} = \hat{\lambda}_{\text{PRS},\text{train}} * \text{PRS}_{\text{test},p}$ |
| 7. Compare the predicted phenotype value to the true phenotype value | $AUC ( \hat{Y}_{\text{test}}, Y_{\text{test}} )$ |

The data was split into the training and testing datasets, with the testing dataset being held out for evaluation. Using the training data, all SNPs’ marginal association, 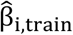, with the biased phenotypes were obtained. The marginal associations from the training data were then used to construct PRSs in both the training and testing data. Next, a predictive model was built using the PRS in the training data; the model was subsequently applied to the testing PRS to obtain the predicted phenotype. The predicted phenotype was compared with the true phenotype in the testing data to obtain the testing area-under-the-curve (AUC) value. The entire process, from phenotype simulation to PRS prediction, was repeated 100 times using different random seeds to obtain 100 replications of the results.

## 3. Result

In all simulations, gold standard results were included to serve as the baselines for comparison. The gold standards demonstrated the maximum obtainable prediction accuracies from PRSs that were generated using the true phenotype. Figure 1 showed a change in PRS prediction accuracy as more non-differential errors were added into the phenotype. The accuracies gradually decreased from gold standard to 50% sensitivity and specificity. At 50% sensitivity and specificity, the biased phenotype was generated the same way as coin-flipping. Thus, the prediction accuracies of PRS at this error level was also around 50%. Notably, the gold standard accuracies were also different even when the simulation parameter values were the same for all three phenotypes.

For differentially misclassified phenotypes, the PRS prediction accuracies also decreased as more differential errors were added to the phenotypes (Figure 3). However, the rates of the accuracy decrease were different for the three types of phenotypes. The PRS derived from the *strongly correlated model* showed the fastest reduction in prediction accuracy. The accuracy from the *weakly correlated model* also decreased faster than that of the *independent model*.

**Figure 2.**
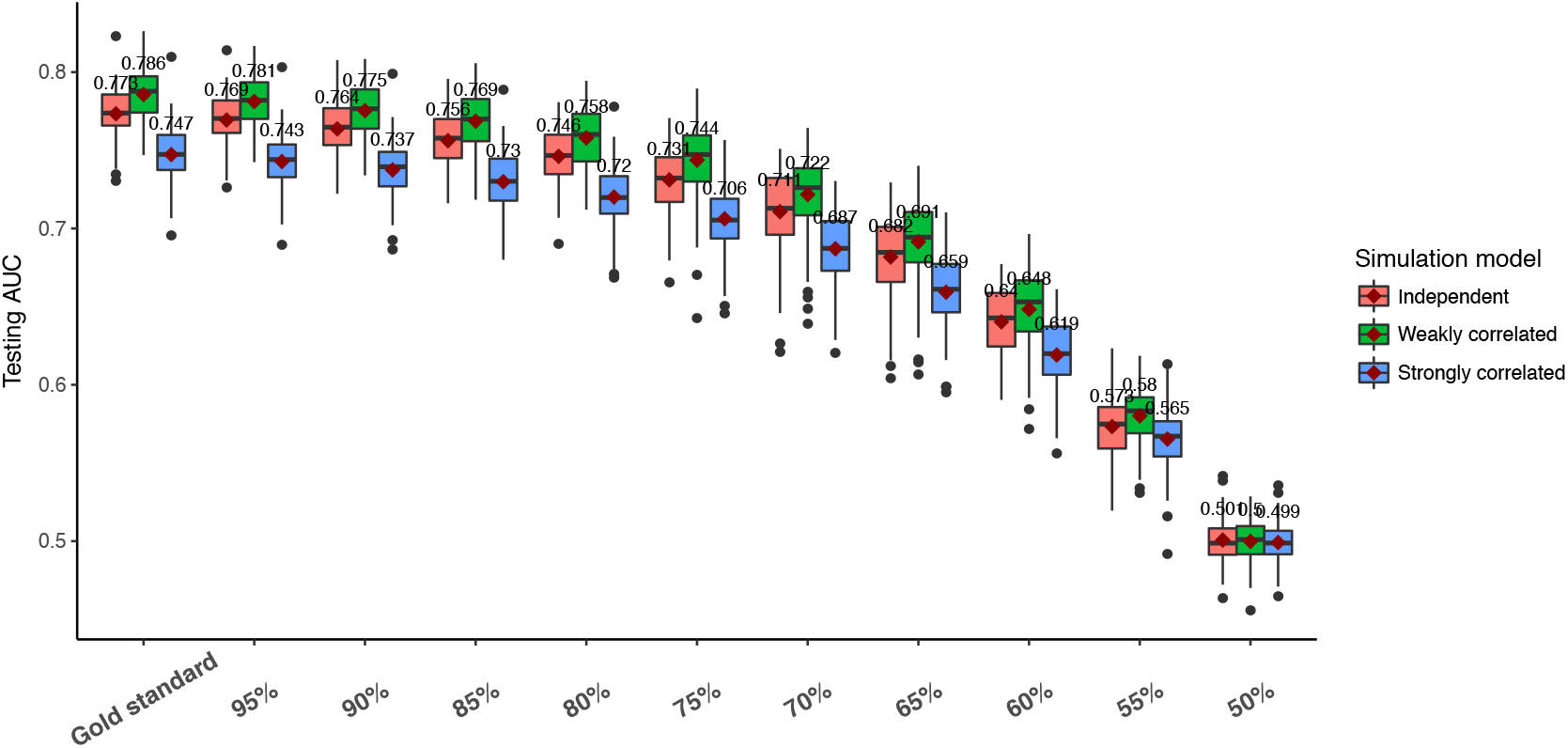
Performance of PRS prediction under non-differential phenotyping error. Each boxplot represents 100 replications of the same experiment using different datasets. The x-axis indicates the sensitivity and specificity level set by variable X in table 2. The y-axis shows the prediction AUC on the testing data.

**Figure 3.**
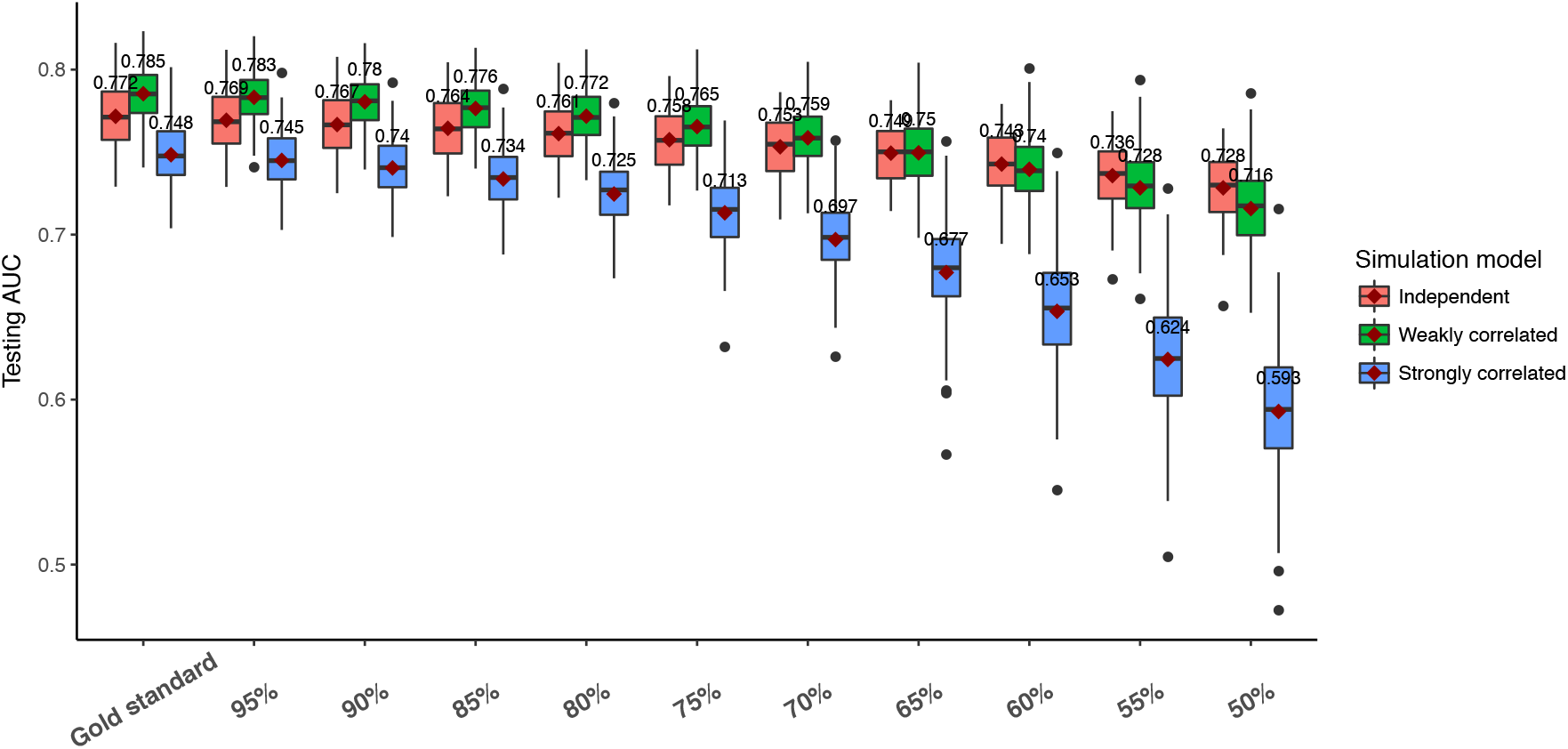
Performance of PRS prediction under differential phenotyping error. Each boxplot represents 100 replications of the same experiment using different datasets. The x-axis indicates the sensitivity and specificity level set by variable X in table 3. The y-axis shows the prediction AUC on the testing data.

## 4. Discussion

Disease risk prediction utilizing genetic information via PRS has shown great promise in many complex human diseases. With the increasing availability of linked genetic data in EHR systems, PRS prediction can be widely applied to many phenotypes and diseases to identify high-risk patients for better disease prevention and treatment care. Nevertheless, patients’ true disease statuses are often unknown. Thus, the observed disease status is only a proxy for the true disease status, and the observed status will be biased due to phenotyping errors. In this study, we quantified the degradation of PRS prediction using three different types of phenotype under the differential and non-differential phenotyping errors.

We utilized the eMERGE EHR genetic data so that the SNPs had the minor allele frequency distribution and correlation structure that are observed in the real patients’ data. Using the SNPs data along with other demographic and clinical variables, we simulated three different phenotypes with increasing levels of complexity (Figure 1). For the phenotype generated under the *independent model*, all variables independently related to the phenotype. Here, we assumed that an individual’s genetic factors do not affect one’s environmental exposure. Under the *weakly correlated* model, we used a related diagnosis status to determine the phenotype status, and the two were associated with a common subset of SNPs through pleiotropic effects. In this case, we assumed that the associated effects were different between the related diagnosis and the phenotype. This is likely when the phenotypes are regulated through different biological mechanisms, such as between heart diseases and mental disorders^22–24^. Finally, in the *strongly correlated* model, the diagnosis and the phenotype were assumed to be more similar due to the shared underlying SNPs as well as their coefficients. This reflects a possible scenario when a subtype of disease is used to diagnose the main disease.

As expected, as more phenotyping errors were added to the three phenotypes, the prediction accuracy of PRS decreased. However, the rates of the decrease depended on the type of phenotyping errors. First, the gold standards’ accuracy in Figure 2 and Figure 3 were similar because they both represented PRS predictive power without any phenotyping errors. Interestingly, the PRS achieved the best performance in the phenotype generated from the *weakly correlated* model, followed by the *independent* and *strongly correlated* model. This can be explained by the different amount of genetic contribution to the phenotype. In the *weakly correlated* model, SNPs contributed to the phenotype through two mechanisms: 1. direct associations with the phenotype. 2. Indirect associations through the related diagnosis. Because the indirect associations were independent of the direct associations, the SNPs contributed “twice” to the phenotype. In contrast, in the *independent model*, the SNPs were only associated with the phenotype through their direct associations. And in the *strongly correlated* model, the SNPs’ associations were diminished because part of the associations was mediated by the related diagnosis. Second, non-differential phenotyping errors similarly affected all phenotypes. The relative order of PRS prediction accuracies did not change as more non-differential phenotyping errors were added. Finally, differential phenotyping errors, which are more likely to be observed in real data, exhibited different accuracy trajectories for the phenotypes. The *independent model* was affected the least, likely because the SNPs and the environmental factor were independent. Thus, differential phenotyping errors induced by the environmental factor did not have a major impact on the PRS prediction accuracy. However, in the *weakly correlated* and *strongly correlated model*, both the phenotype and the related diagnosis were associated with the SNPs. Thus, differential errors based on these variables had a severe impact on the PRS, with the strongest impact in the *strongly correlated model.* In summary, non-differential phenotyping errors affected PRS prediction equally among the phenotypes. Differential phenotyping errors had an increased impact on PRS prediction if the target phenotype and the variables used to determine the phenotype have a shared genetic component.

While it is useful to understand the impact of phenotyping errors on PRS prediction, it is also important to identify approaches that can minimize the error. One effective approach to reducing error is through manual chart review of patients’ comprehensive clinical histories by doctors or domain experts. However, manual review is both time-consuming and expensive. A potential alternative approach is to chart review a subset of patients to determine the amount of phenotyping error as well as the error mechanism. Then, the results presented in this study could serve as a guideline to determine whether the errors are within the acceptable range. If not, the phenotype quality needs to be improved. For future studies, the impact of phenotyping errors on the continuous outcome can be explored. Furthermore, more complex error patterns that depend on multiple environmental or clinical variables are likely to be more realistic and should be investigated. Finally, some studies suggested that AUC may not be the best metric for evaluating classification accuracy. Thus, other accuracy metrics, such as net reclassification improvement or integrated discrimination improvement can be used^25^.

## Notes

* This work is supported by 1R01LM012607, 1R01AI130460, and LM010098.

